# *Actinobacillus* utilizes a binding-protein dependent ABC transporter to acquire Vitamin B_6_

**DOI:** 10.1101/2020.05.01.072959

**Authors:** Chuxi Pan, Alexandra Zimmer, Megha Shah, Minhsang Huynh, Christine C.L. Lai, Brandon Sit, Yogesh Hooda, David Curran, Trevor F. Moraes

## Abstract

Bacteria require high efficiency uptake systems to survive and proliferate in nutrient limiting environments, such as those found in the host. The ABC transporters at the bacterial plasma membrane provide a mechanism for transport of many substrates. We recently demonstrated that an AfuABC operon, previously annotated as encoding a ferrous iron uptake system, is in fact a cyclic hexose/heptose-phosphate transporter with high selectivity and specificity for these metabolites. In this study, we examine a second operon containing a periplasmic binding protein discovered in *Actinobacillus* for its potential role in nutrient acquisition. Using electron density obtained from the crystal structure of the periplasmic binding protein we modeled a pyridoxal-5’-phosphate (P5P/PLP/Vitamin B_6_) ligand into the atomic resolution electron density map. The identity of the Vitamin B_6_ bound to this periplasmic binding protein was verified by isothermal titration calorimetry, microscale thermophoresis, and mass spectrometry, leading us to name the protein P5PA and the operon P5PAB. To illustrate the functional utility of this uptake system, we introduced the P5PAB operon from *A. pleuropneumoniae* into an *E. coli* K-12 strain that was devoid of a key enzyme required for Vitamin B_6_ synthesis. The growth of this strain at low levels of Vitamin B_6_ supports the role of this newly identify operon in Vitamin B_6_ uptake.

## Introduction

*Actinobacillus pleuropneumoniae* is a Gram-negative bacterial pathogen that preferentially infects and colonizes the lower respiratory tracts of pigs. It is the causative pathogen for the highly contagious porcine pleuropneumonia which is characterized by pulmonary edema, hemorrhage and necrosis(1, 2). Acute outbreaks of *A. pleuropneumoniae* infection can result in swine death within 24 hours, leading to high economic losses for the swine industry(1, 3). Antibiotic therapy is the most common and effective means of restricting the spread of infection and decreasing mortality. However, *A. pleuropneumoniae* strains with antibiotic resistance have been recently reported(1). Therefore, it is important to understand the pathogenesis of *A. pleuropneumoniae* and develop alternative treatments. Previous studies have identified virulence factors used by *A. pleuropneumoniae* to acquire essential nutrients (2). These are required for successful host invasion but are either sequestered or not available in the respiratory tract(2). Thus, efficient nutrient acquisition systems, such as binding protein dependent transporters (BPDTs) have been evolved in *A. pleuropneumoniae* and provide it with the necessary nutrients to outgrow competing organisms.

BPDT systems are composed of three components: a periplasmic binding protein and a permease/ATPase complex embedded in the inner membrane. The periplasmic binding protein is able to bind to a specific substrate and recruits it to the permease/ATPase complex, which then uses ATP hydrolysis to induce substrate influx from the periplasm into the cytosol(4). Despite playing an important role in nutrient acquisition, BPDT systems in *A. pleuropneumoniae* have been either misannotated or gone completely uncharacterized. For example, the BPDT system designated AfuABC (*Actinobacillus* ferric uptake ABC) was identified in *A. pleuropneumoniae* and, based on sequence homology to FbpA, the periplasmic binding protein AfuA was proposed to be a transporter for ferric iron(5). However, the crystal structure of AfuA in complex with glucose-6-phosphate (G6P) revealed that AfuABC system is a highly specific transporter for hexose/heptose-phosphate molecules and provides a significant alternative carbon source for bacterial metabolism in the lower gut(6). Following the identification of the role of AfuA, we discovered another gene (APJL_RS02700) in *A. pleuropneumoniae* with high sequence identity to AfuA, and began to characterize the unknown structure and function of this putative periplasmic binding protein.

In this study, we solve the structure of the periplasmic binding protein encoded by APJL_RS02700 in complex with a ligand, and identify that ligand to be pyridoxal-5’-phosphate (P5P/PLP/Vitamin B_6_). We confirm the identity of the ligand in the binding pocket using mass spectrometry, and further show, using isothermal titration calorimetry (ITC) and microscale thermophoresis (MST), that Vitamin B_6_ binds our periplasmic binding protein. Lastly, we demonstrate that expressing the operon containing APJL_RS02700 in a strain of *E. coli* deficient in Vitamin B_6_ synthesis rescues its growth in media containing low levels of P5P. We propose the name P5PA, for Pyridoxal 5 Phosphate binding protein A or Vb6A, for Vitamin B_6_ binding protein A, for this newly characterized periplasmic binding protein. To our knowledge, P5PA is the only bacterial protein shown to promote uptake of the active form of vitamin B_6_.

## Materials and methods

### Bacterial strains and growth conditions

All bacterial strains used in this study are listed in Table S1. All *Escherichia coli* strains were grown in Luria-Bertani (LB) broth or M9 minimal media (Bioshop) or on LB agar plates at 37°C. When antibiotics were needed for plasmid selection, they were added at the following concentrations: 50 μg/mL kanamycin and 50 μg/mL ampicillin.

### Cloning and expression of P5PA, P5PB (APJL_RS02705) and AfuC

Plasmids used in this study are listed in Table S1. To construct pET26b-HisP5PA, the coding sequence of P5PA was amplified from the genomic DNA of *A. pleuropneumoniae* strain H49 and integrated into a pET26b expression vector by restriction free cloning(7). To construct the plasmids pSC101-P5PAB and pSC101-P5PAB+AfuC, the operon containing *p5pAB* was cloned into the low-copy customized vectors pSC101 and pSC101-AfuABC by exponential megaprimer cloning(8). pSC101-AfuABC was obtained from the previous study performed by Sit et al.(6). All constructed plasmids were verified by TCAG DNA sequencing (The Centre for Applied Genomics, Toronto, Canada).

### P5PA protein purification

The plasmid pET26b-HisP5PA was transformed into *E. coli* BL21 competent cells which were then grown overnight in LB broth supplemented with 50 μg/mL kanamycin at 37°C with shaking. The overnight culture was then used to inoculate 2YT media (1:100) for large-scale protein expression. IPTG was added to induce protein overexpression with a concentration of 0.5 mM when OD_600_ reached 0.6. After 4-hours of incubation at 20°C, cells were harvested by centrifugation at 7500 rpm at 4°C and resuspended in Lysis buffer (10 mM imidazole, 50 mM Tris pH 8 and 300 mM NaCl) containing 1 mM phenylmethylsulfonyl fluoride (PMSF), 1 mM benzamidine, 20 mg/mL lysozyme and 5 mg/mL DNases. The cell suspension was sonicated for 15 minutes with 15s ON/30s OFF intervals to lyse the cells This was followed by centrifugation at 17,000 rpm for 30 minutes to pellet cell debris(9). The supernatant was passed through a 0.45 μm syringe filter, combined with 1mL of Ni-NTA resin, and incubated at 4°C for at least one hour. The solution was then loaded into a gravity column, washed with Lysis buffer and eluted with Elution buffer (400 mM imidazole, 50 mM Tris pH 8 and 300 mM NaCl). Subsequently, the eluted protein was dialyzed with thrombin in the Lysis buffer to cleave off 6xHis-tag from the N-terminus of P5PA. After overnight dialysis at 4°C, Ni-NTA resin and benzamidine resin were used to remove uncleaved His-P5PA, thrombin and the free 6xHis-tag. The protein solution was then centrifuged at 13,000 rpm for 10 minutes and concentrated in a centrifugal concentrator with a 10 kDa cutoff (Millipore, USA). Finally, the protein was loaded onto a Superdex 75 10/300 gel filtration column (GE Healthcare, USA), which had been equilibrated in buffer containing 20 mM Tris pH 8.0 and 100mM NaCl, for further purification(6). The purity and yield of protein was estimated by SDS-PAGE and 280 nm readings made on a Nanodrop ™, respectively. The N-terminal 6xHis-tag was only cleaved for P5PA used in X-ray crystallography trials.

### Crystallization, data collection and structure determination

P5PA was concentrated to 12 mg/mL and used in screen-based crystallization trials. The Gryphon robot (Art Robbins Instrument) was used to set up the MSCG 1-4, JCSG1 and Index commercial screens. P5PA was initially crystallized in two hit conditions: (1) 0.1 M potassium thiocyanate, 30% PEG MME 2000; (2) 0.1 M CHES: NaOH pH 9.5, 30% PEG 3000. The crystal hits were manually optimized by varying the pH and concentrations of polyethylene glycol (PEG) and glycerol. Candidate crystals were looped into a cryoprotectant solution consisting of the mother liquor with 20% glycerol and flash frozen in liquid nitrogen. The crystals were sent to the Advanced Photon Source (APS) Synchrotron Facility (Argonne, USA) for data collection. Molecular replacement (MR) was performed by running Phaser in PHENIX(10) using AfuA (PDB ID: 4R73) as the template (54% sequence identity shared with P5PA). MR and automated refinement were performed using PHENIX. Manual refinement was performed using Coot(11). Molecular models (Fig. 3) were generated using PyMol (Warren De Lano).

### Isothermal Titration Calorimetry (ITC)

The purification method described in the **P5PA protein purification** section was modified as follows in order to purify apo-P5PA for ITC. Before elution, P5PAwas denatured in the gravity column. To denature P5PA, 6 M guanidine hydrochloride (GdHCl) was applied to the column, after which a refolding step was performed by stepwise addition of 4.5 M GdHCl, 3 M GdHCl, and finally 1.5 M GdHCl. The protein was then washed with Lysis buffer and eluted with Elution buffer(6). The subsequent purification steps were the same as previously described.

ITC was performed using an ITC-200 instrument (Structure & Biophysical Core Facility, Peter Gilan Centre for Research & Learning, Canada). Runs consisted of titrating ligand loaded into the injection syringe against purified protein, which was loaded into the sample cell. 45 μM of purified P5PA was stored in ITC buffer (20 mM Tris pH 8 and 100 mM NaCl) at 4°C. The same buffer was used to make solutions of 300 μM glucose-6-phosphate(G6P), 300 μM P5P, ands 45 μM bovine serum albumin (BSA). P5PA, BSA and ITC buffer alone were titrated with Vitamin B_6_ or G6P. The parameter setting was 2 μL/injection for a total of 20 injections. Data applied to calculate binding constants were referenced against runs performed with ligand alone to control for the heat of ligand solvation.

### Identification of the P5PA ligand by tandem liquid chromatography/mass spectrometry

Endogenous ligands of P5PA were sought by liquid-liquid extraction. 1 mL of 80% cold acetone (−20°C) and 20% cold 2 M HCl was added to P5PA in ITC buffer. The mixture was then vortexed for 5 minutes, incubated at 4°C overnight and centrifuged at 15,000 rpm for 10 minutes. The supernatant was harvested and dried by SpeedVac SC100 (Savant). The dried pellet was dissolved in milliQ water, and the solution was then used for liquid-liquid extraction with acetonitrile and saturated NH4Cl to remove excess NaCl and Tris from the solution(12, 13). The organic layer was transferred to a new tube and dried. The dried precipitates were dissolved in milliQ water and sent to the Analytical Facility for Bioactive Molecules (Hospital for Sick Children, Canada) for tandem liquid chromatography/mass spectrometry (LC-MS/MS).

### *E. coli* growth assays

The *ΔpdxB* mutant of *E.coli* K-12 strain was obtained from the Keio collection(14). The *ΔpdxB* mutant was transformed with either pSC101, pSC101-P5PAB, or pSC101-P5PAB+AfuC and grown in LB broth overnight at 37°C with shaking. The starters were then washed twice with M9 medium and inoculated at OD_600_=0.05 into M9 medium supplemented with Vitamin B_6_. The Vitamin B_6_ concentrations used were 0 μM, 1μM, 2.5 μM and 5 μM. Growth assays were performed in both culture tubes and 100-well microplates (6, 15). For cell growth in culture tubes, 5mL of P5P-supplemented M9 was used and OD_600_ readings were measured after a 23-hour incubation at 37°C with shaking. For cell growth in 100-well microplates, 200 μL of P5P-supplemented M9 was applied to each well and microplates were incubated in a Bioscreen C microplate reader (Growth Curves USA) at 37°C with shaking. OD_600_ values were measured every 15 minutes for a time course of 40 hours.

### *In silico* detection of P5PA

As P5PA and AfuA share significant sequence similarity (54% amino acid identity), a reciprocal BLAST approach was used to ensure that we did not mistakenly identify any AfuA as P5PA. We first queried a P5PA protein sequence (accession WP_012262855) against the nr database using BLASTp (v2.10.0+)(16). The top 100 hits shared >66% identity, so an additional seven sequences were chosen as a set of “seed sequences” for which we had high confidence in their function. These 8 seed sequences were then queried against the refseq protein database using BLASTp, excluding hits from all 8 seed species. This returned 64 hits that shared >65% identity over >80% of the sequence with at least one seed sequence. Nine hits showed similarity to only one or two seed sequences, and so were added to the set of seed sequences to improve our detection ability. These 17 seed sequences were then queried against the refseq_protein database using BLASTp, again excluding all hits from the seed species. This returned 88 hits sharing >65% identity over >80% of the sequence.

A similar process was used to identify eight AfuA seed sequences, which were queried against the refseq protein database using BLASTp, and returned 234 hits sharing >65% identity over >80% of the sequence. The 88 putative P5PA hits were then queried against the set consisting of 17 P5PA seed sequences, 8 AfuA seed sequences, and 234 AfuA sequences using BLASTp. P5PA hits were only considered valid if all three of the most similar sequences were from the P5PA seed sequences, not from the set of AfuA sequences. All 88 hits passed this threshold, and so when combined with the seed sequences we were able to identify a total of 105 P5PA sequences.

## Results

### The structure of P5PA

Initial screening identified two nucleation conditions: 0.1 M potassium thiocyanate, 30% PEG MME 2000 and 0.1 M CHES:NaOH pH 9.5, 30% PEG 3000. These conditions were manually optimized to produce crystals, which diffracted at a resolution of 1.75 Å at the APS (see Methods). Using molecular replacement with AfuA as a model, we solved the structure of P5PA to a resolution of 1.75 Å (see Methods). The resulting high-resolution structure revealed that P5PA is a type-II periplasmic binding protein with two β-strands connecting two globular domains. The P5PA binding cleft is located above the β-strands at the interface of the two globular domains (Fig. 1A) and was observed to have positive density (Fig. 1B). A screen for possible ligands revealed that neither glucose-6-phosphate nor fructose-6-phosphate, the two primary ligands that bid AfuA, fit the positive density in the binding pocket. Further screening of potential ligands containing six-member rings and phosphate groups identified the active form of vitamin B_6_, called pyridoxal-5’-phosphate (P5P/PLP), as the best fit (Fig. 1A).

**Figure 1.**
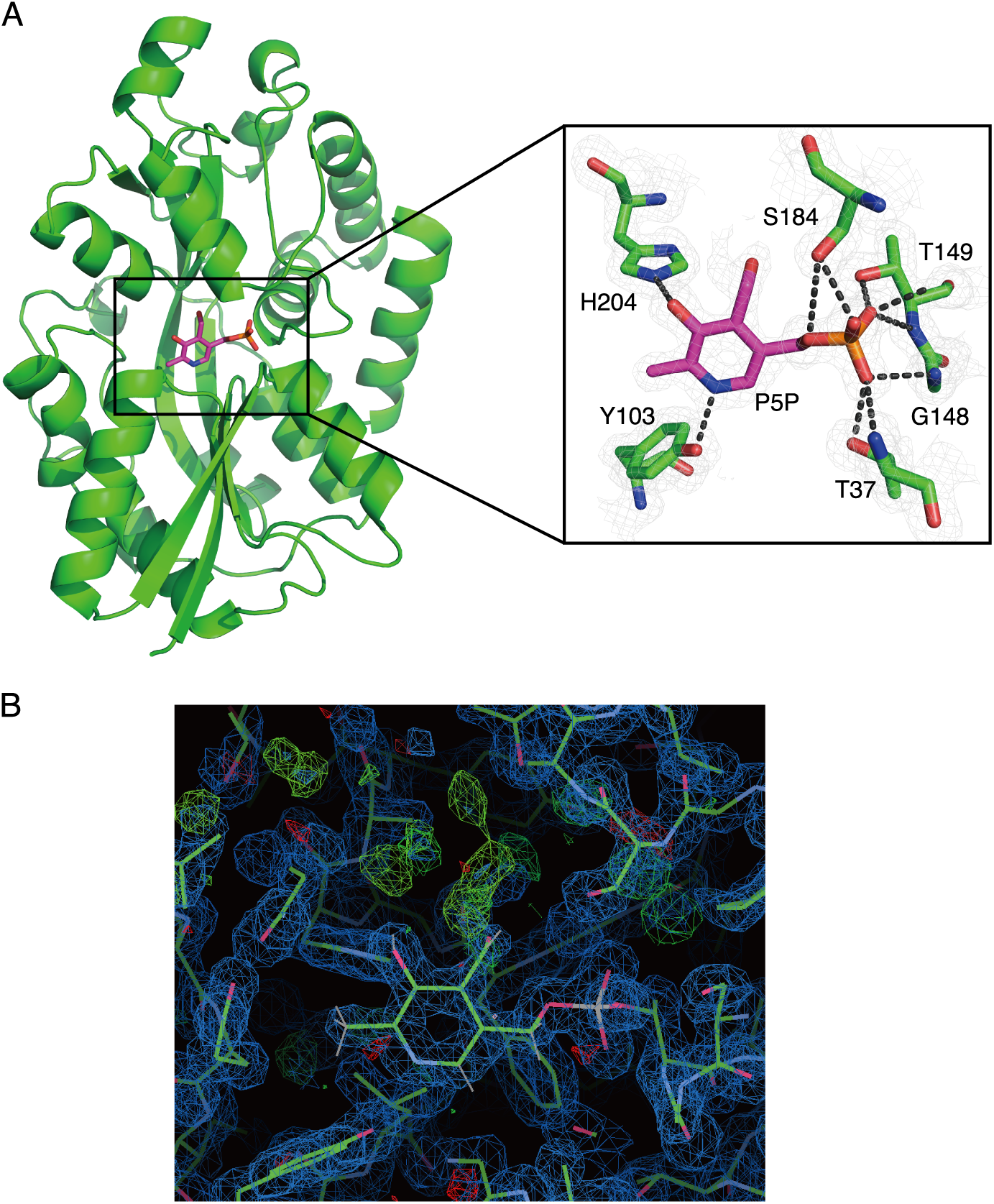
P5PA is a potential transporter for Vitamin B_6_. **(A)** Ribbon diagram of P5PA (green) bound to endogenous Vitamin B_6_ (P5P magenta) (PDB code: 6WCE determined in this study). The co-purified Vitamin B_6_ is shown in the inset with interacting residues shown as sticks. The pyrimidine ring of P5P (Vitamin B6) is coordinated by Y103 and H204 while the phosphate group are stabilized together by T37, G148, T149 and S184. **(B)** Electron density map of P5PA with Vitamin B_6_ (P5P) modelled in the binding pocket.

The model of P5PA with Vitamin B_6_ in the binding pocket reveals a network of hydrogen bonds holding the pyrimidine ring and phosphate group of Vitamin B_6_ in place. While Tyr103 interacts with the pyrimidine ring nitrogen and His204 coordinates the pyrimidine hydroxyl group, the Vitamin B_6_ phosphate group is stabilized by interactions with the side chains of Thr37, Ser183, and Thr149, and the backbone of Gly148 (Fig. 1A). Interestingly, after we refined the structure with the Vitamin B_6_ ligand in the binding site, we found unmodelled density at carbon 4 of P5P (Fig. 1B), indicating that derivatives of Vitamin B_6_ are also likely to be ligands of P5PA.

### Measuring the binding affinity between P5PA and Vitamin B_6_ using ITC

Because the structure of the P5PA-Vitamin B_6_ complex showed additional density near carbon 4 of Vitamin B_6_, we sought to determine whether Vitamin B_6_ is truly a ligand of P5PA. To do this, we used ITC to measure the binding affinity between these two molecules. We determined a dissociation constant (K_d_) of 35.53 +/- 7.33 nM, which suggests that there is a strong interaction between P5PA and Vitamin B_6_ (Table 1; Fig. 2A). We detected no interaction between P5PA and bovine serum albumin (BSA), which functioned as a negative control (Fig. 2B).

**Table 1.**
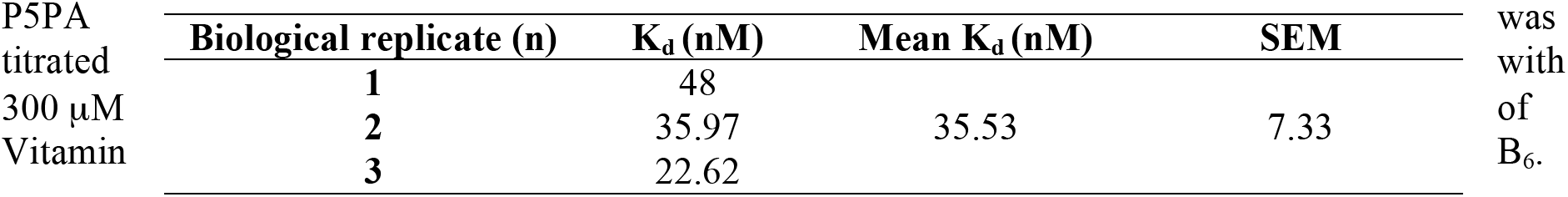
Dissociation binding constants (K_d_) for the interaction between P5PA and Vitamin B_6_. 45μM of Released Heat, K_d_ and stoichiometries were measured and analyzed by ITC-200. Biological replicates n=3 were completed.

| P5PA<br>titrated<br>300 $\mu\text{M}$<br>Vitamin | Biological replicate (n) | $K_d$ (nM) | Mean $K_d$ (nM) | SEM | was<br>with<br>of<br>B<sub>6</sub>. |
| --- | --- | --- | --- | --- | --- |
|  | 1 | 48 |  |  |  |
|  | 2 | 35.97 | 35.53 | 7.33 |  |
|  | 3 | 22.62 |  |  |  |

**Figure 2.**
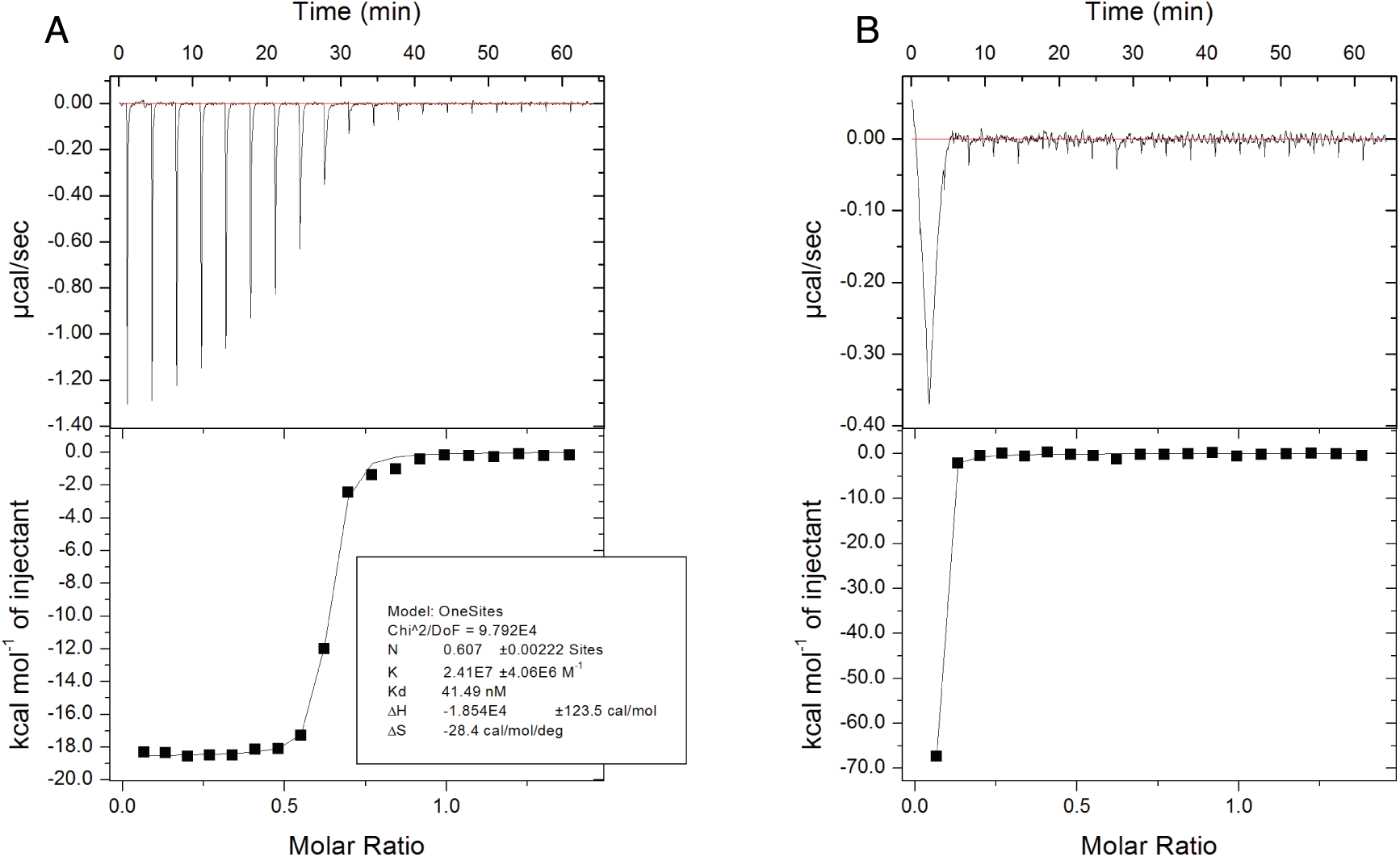
P5PA has a strong and specific interaction with Vitamin B6. **(A)** Representative ITC data showing the interaction between P5PA and Vitamin B_6_. 45 μM of P5PA was titrated with 300 μM of Vitamin B_6_. Heat release was measured by ITC-200. **(B)** ITC data showing the interaction between BSA and Vitamin B_6_ (negative control). 45μM of BSA was titrated with 300 μM of Vitamin B_6_. The other parameters were the same as **(A)**.

As P5PA is a homologue of AfuA, we wondered whether P5PA is able to interact with hexose phosphate molecules. We measured the interaction between glucose-6-phosphate (G6P) and P5PA using ITC. The released heat was approximately 3 kJ/mol for each G6P to P5PA titration, which was insufficient to calculate a dissociation constant (Fig. S1). This suggests that there is no significant binding activity between G6P and the periplasmic binding protein P5PA. Taken together, we conclude that P5PA specifically binds its ligand Vitamin B_6_ with high affinity.

### Identification of P5PA by mass spectrometry

To confirm that Vitamin B_6_ is a ligand of P5PA and to determine if P5PA has any other potential substrates, we extracted ligands from a solution of ligand-bound P5PA and performed liquid chromatography/tandem mass spectrometry (LC-MS/MS). The molecular weight of Vitamin B_6_ (P5P) is 247 Da. As a proton was added to ionize Vitamin B_6_, the mass-to-charge ratio (m/z) representing Vitamin B_6_ became 248. We used Vitamin B_6_ powder dissolved in ultrapure water as a positive control. The peak representing 248 m/z was detected in both the positive control Vitamin B_6_ solution and the ligand extract, although the peak in the Vitamin B_6_ solution had higher intensity (Table 2; Fig. 3A; Fig. 3C). In order to identify the molecule represented by the detected 248 m/z peak, the products at 248 m/z were fragmented by nitrogen gas. The expected transition of 248 → 150 m/z can be used to characterize Vitamin B_6_ molecules(17). The 248 → 150 m/z fragmentation pattern was observed in both the extract (Fig. 3D) and P5P solution (Fig. 3B), implying that Vitamin B_6_ was present in the extract of ligand-bound P5PA. Overall, the major peaks present in the Vitamin B_6_ solution and the extract had the same mass-to-charge ratios but different intensities, suggesting that no other Vitamin B_6_ derivatives were identified by LC-MS/MS.

**Table 2.** LC-MS analysis of ligands extracted from holo-P5PA and dissolved Vitamin B_6_ solution.

|  | Original concentration (μM) | 248 m/z intensity |
| --- | --- | --- |
| Vitamin B<sub>6</sub> solution | 248 | 9x10<sup>6</sup> |
| Substrates extracted from holo-P5PA | 248 | 7.8x10<sup>6</sup> |

**Figure 3.**
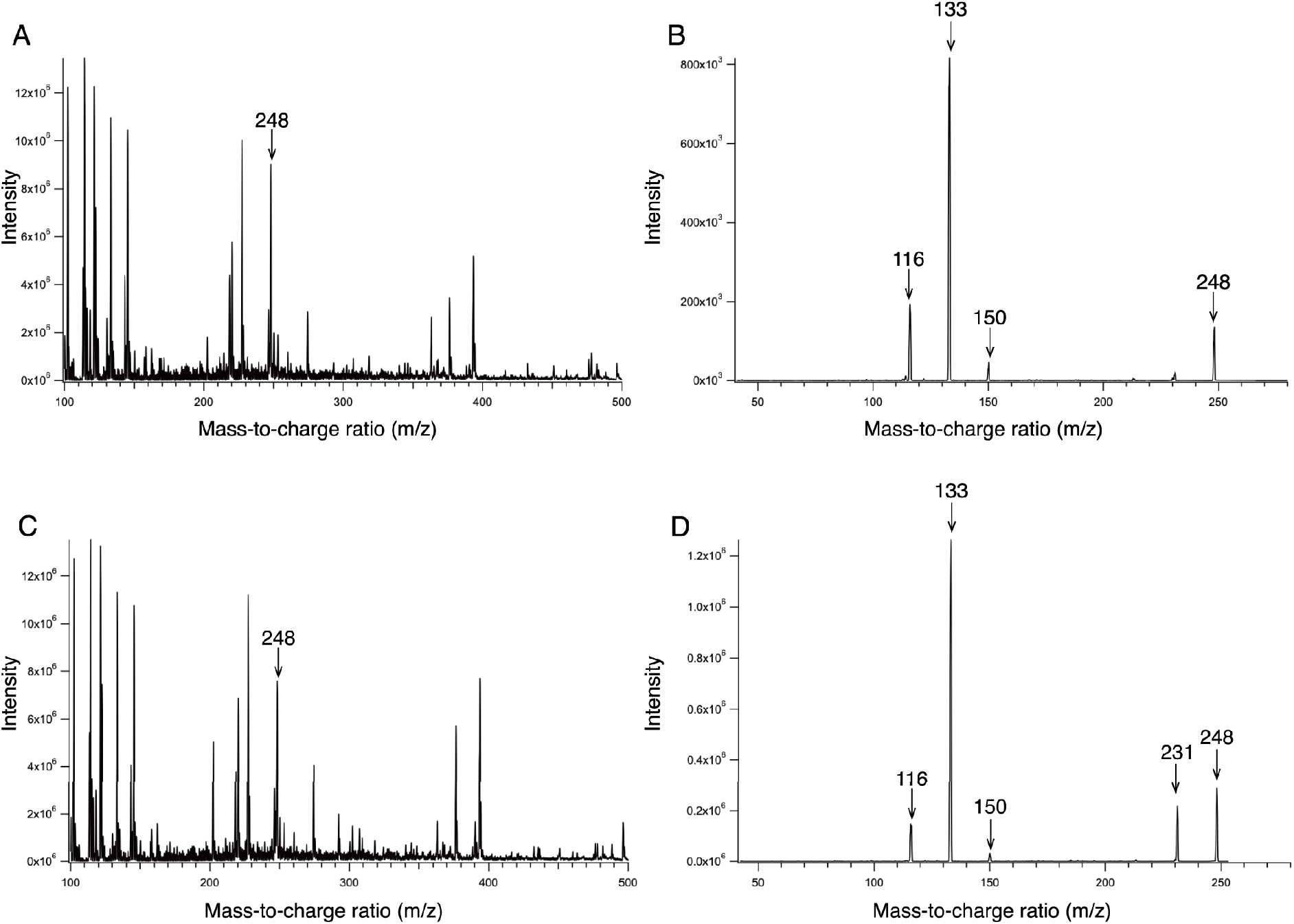
LC-MS/MS profiles of Vitamin B_6_ solution and holo-P5PA extract. Holo-P5PA was purified and added to acidic acetone to remove protein. The extract, which contained the endogenous ligand, was harvested. Acetonitrile and saturated NH4Cl were applied to remove Tris and NaCl and the extract was then was then subjected to tandem liquid-liquid chromatography/mass spectrometry. Dissolved Vitamin B_6_ (P5P) solution was used as a positive control. The presence of Vitamin B_6_ was indicated by a peak at 248 m/z. (A) LC-MS/MS profile of the Vitamin B_6_ solution from 100-500 m/z. (B) The fragmented Vitamin B_6_ solution peak at 248 m/z. (C) LC-MS/MS profile of the extract from 100-500 m/z. (D) The fragmented extract peak at 248 m/z.

### P5PA facilitates *E. coli* growth at low levels of Vitamin B_6_

Since our results reveal that Vitamin B_6_ is a ligand of P5PA *in vitro*, we wondered whether binding of P5PA to Vitamin B_6_ was important *in vivo*. To address this, we asked whether expression of the vitamin B_6_ uptake system from *A. pleuropneumoniae* could rescue the growth of bacterial strains deficient in Vitamin B_6_ synthesis. In wild-type *E. coli* (strain K-12), Vitamin B_6_ can be synthesized from D-erythrose 4-phospate, pyruvate and glyceraldehyde 3-phosphate by enzymes encoded by *pdx* family genes (Fig. 4). When the Vitamin B_6_ biosynthesis pathway is disrupted, *E.coliK-12* displays severe growth defects in the absence of exogenous Vitamin B_6_ (15). The *E. coli* K-12 auxotroph, *ΔpdxB*, contains a deletion of 4-phosphoerythronate dehydrogenase that plays a significant role in converting D-erythrose 4-phosphate to Vitamin B_6_.

**Figure 4.**
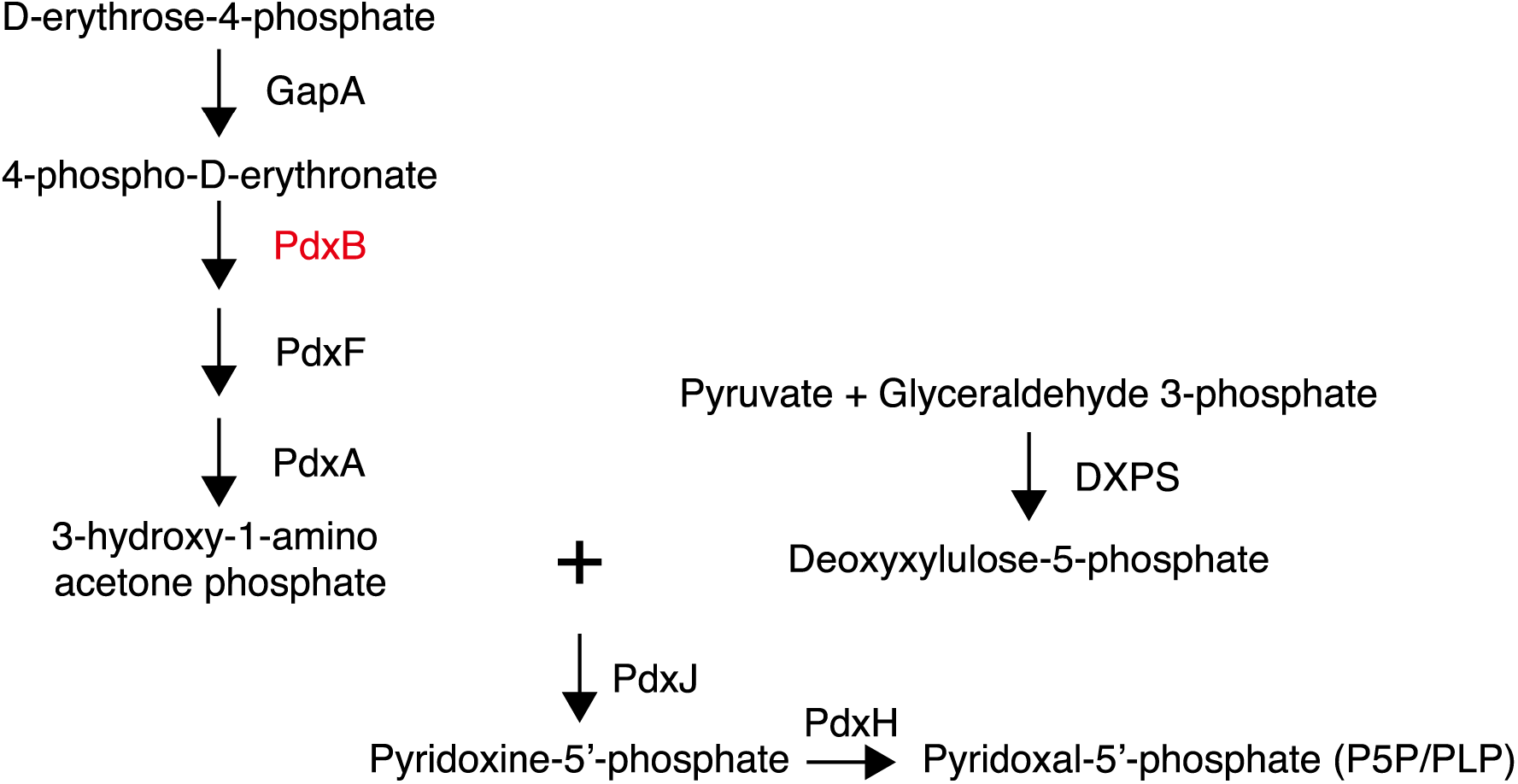
Vitamin B_6_ de novo biosynthesis pathway in *E. coli* K-12. In *E. coli* K-12, D-erythrose-4-phosphate, pyruvate and glyceraldehyde-3-phosphate can be used to synthesize P5P. The synthesis is catalyzed by GapA, DXPS and enzymes encoded by the pdx genes. Each arrow represents an enzymatic reaction between a pathway intermediate and the enzymes alongside the arrows.

We tested K-12::Δ*pdxB* for its growth capacity in M9 minimal media supplemented with Vitamin B_6_ at different concentrations varying from 0 μM to 10 μM. After 24 hours of incubation, we found that the wild-type strain (WT) of *E. coli* K-12 grew to stationary phase and that its growth was not affected by the concentration of supplemented Vitamin B_6_. However, the *ΔpdxB* mutant had severe growth deficiency in all conditions except Vitamin B_6_ concentrations of 5 μM and 10 μM (Fig. S2). As the *ΔpdxB* mutant showed a measurable growth deficiency at low concentrations of P5P, it was used to investigate the role of P5PA.

P5PA is encoded by the *p5pAB* operon in *A. pleuropneumoniae*. BPDT operons typically encode a periplasmic binding protein, transmembrane domains, and ATPases. However, the gene encoding the ATPase component was not present in the downstream of *p5pAB*. Therefore, we have two hypotheses to explain how P5PAB functions. The first hypothesis is that the gene annotated “*p5pB*” encodes both the transmembrane domains and ATPases. The other possibility is that the *p5pAB* operon shares ATPases with other BPDT systems. Other studies have demonstrated that the ATPase component MsiK is shared by a couple of ABC transport systems for disaccharides in Gram-positive bacteria(18, 19). As P5PA shares 54% amino acid sequence identity with its homologue AfuA and P5PB shares 60% identity with AfuB, we believe that P5PAB may be able to utilize the ATPase component, AfuC, from the AfuABC hexose-phosphate uptake system. Thus, we made two constructs, one containing the *p5pAB* operon and another containing the *p5pAB* operon together with *afuC*.

In order to rescue growth of the *ΔpdxB* mutant, this strain was transformed with either pSC101 empty vector, pSC101-P5PAB, pSC101-P5PAB+AfuC, or pSC101-AfuABC and grown in M9 media supplemented with Vitamin B_6_. When Vitamin B_6_ was supplied at concentrations of 1 μM and 2.5 μM, growth of the *ΔpdxB* mutant was rescued by the expression of pSC101-P5PAB+AfuC (p<0.01, two-tailed t-test); however, expression of pSC101, pSC101-P5PAB, and pSC101-AfuABC did not have any effect on the growth of the *ΔpdxB* mutant. When Vitamin B_6_ was at 5 μM, the growth defects of the four transformants and the *ΔpdxB* mutant were partially restored (Fig. 5) while the pSC101-P5PAB+AfuC transformant grew the fastest of all the transformants to the stationary phase (Fig. 6D). Given our results, we conclude that P5PAB on its own does not function efficiently to acquire exogenous Vitamin B_6_ and requires a shared ATPase component from other ABC transport systems. In this case, the ATPase AfuC supplies sufficient energy to allow for Vitamin B_6_ uptake. Moreover, this experiment verifies that P5PA performs Vitamin B_6_ uptake.

**Figure 5.**
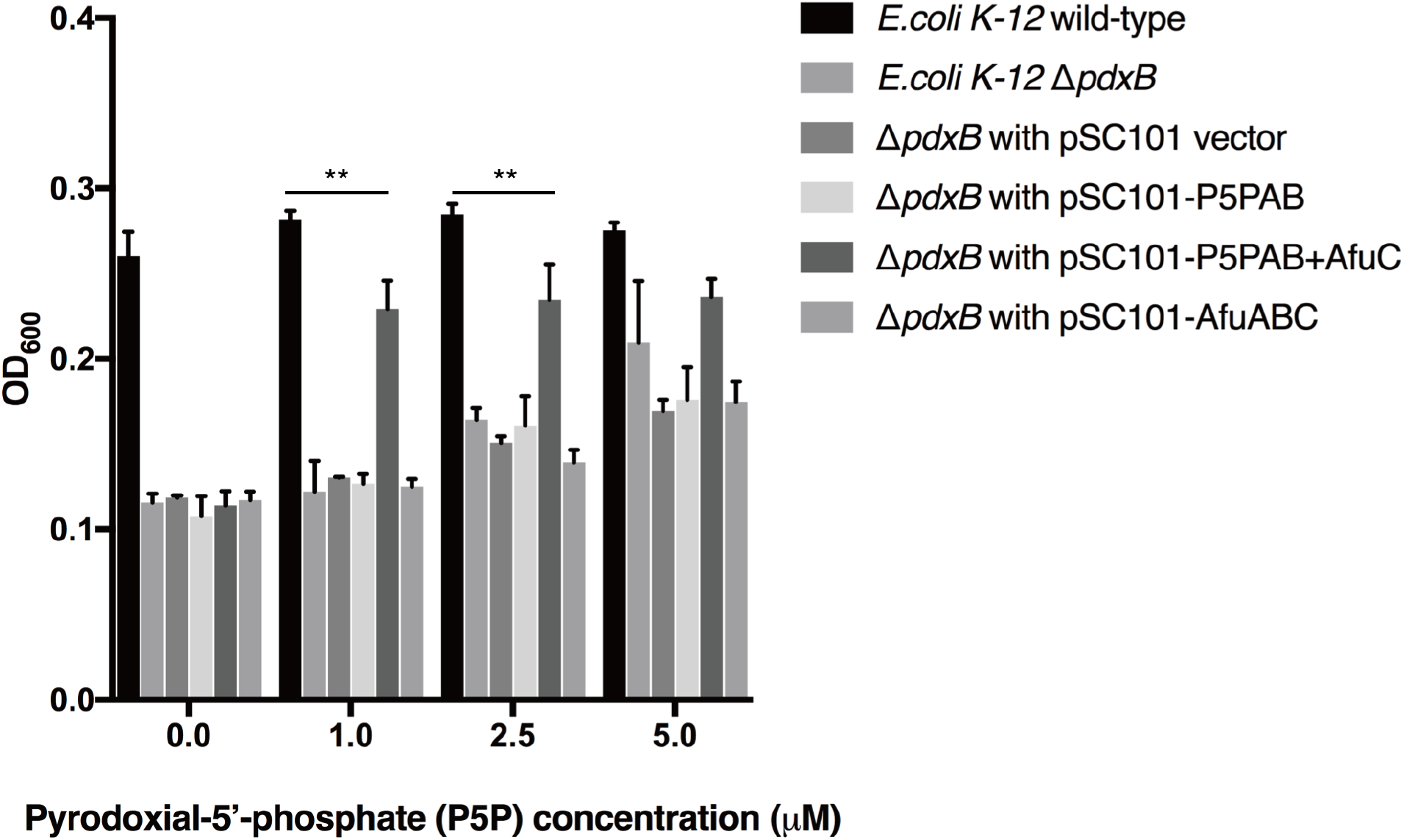
*p5pAB+afuC* rescues growth of the *E. coli* K-12 *ΔpdxB* mutant. *E. coli* K-12 *ΔpdxB* mutants were transformed with four constructs: pSC101, pSC101-P5PAB, pSC101-P5PAB+AfuC and pSC101-AfuABC. They were then incubated in 200 μL of M9 supplemented with Vitamin B_6_ (Pyrodoxial-5’-phosphate –P5P) at 1 μM, 2.5 μM and 5 μM. The starting OD_600_ was 0.08. The bars represent the mean OD_600_ readings after 40 hours. The error bars represent standard deviations from n=3 transformants. (*P<0.05, **P<0.01)

**Figure 6.**
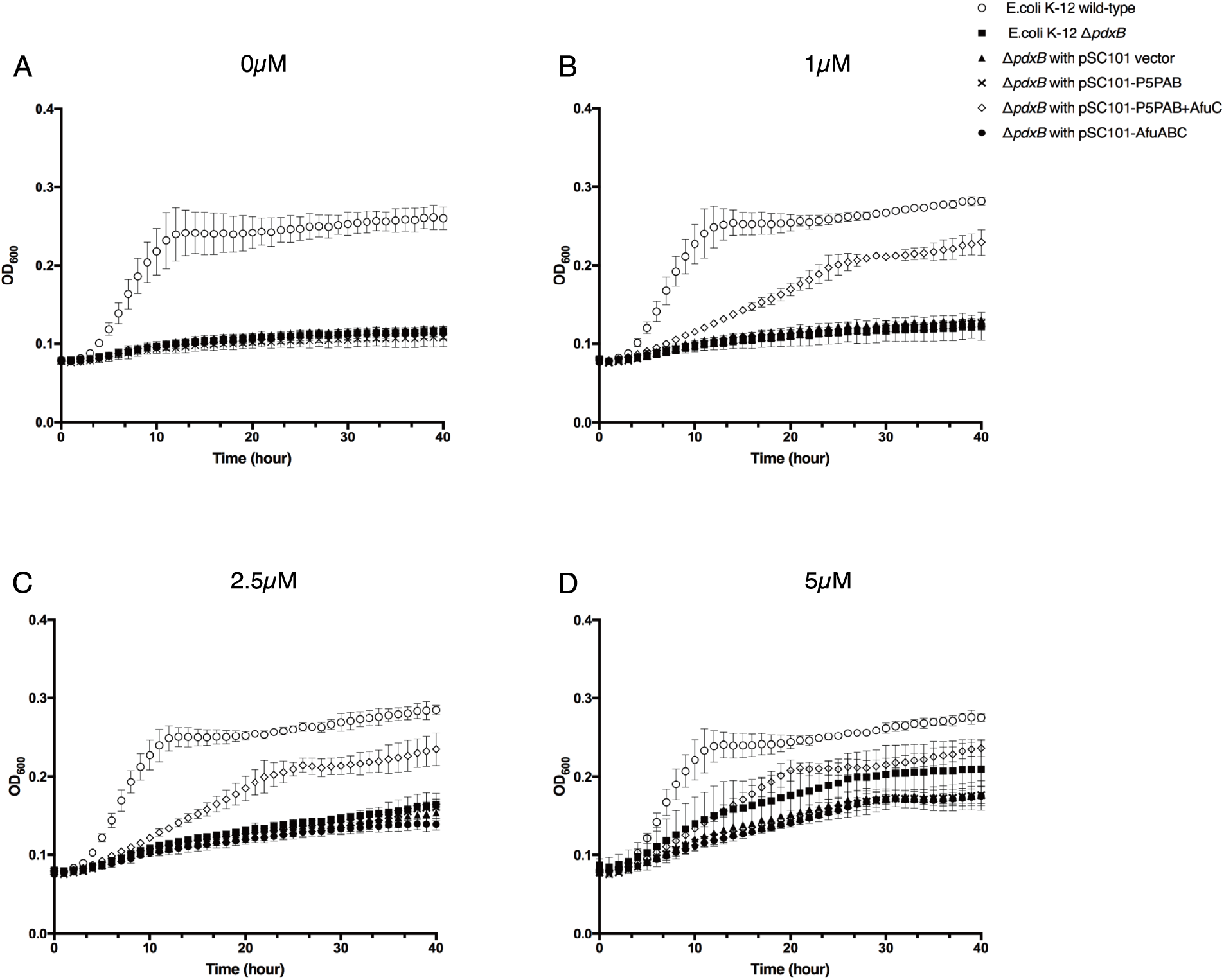
*afuA2B2C* restored the growth rate of *E. coli* K-12 Δ*pdxB* mutant. *E. coli* K-12 Δ*pdxB* mutants were transformed with four constructs: pSC101, pSC101-AfuA2B2, pSC101-AfuA2B2C and pSC101-AfuABC. They were then incubated in 200 μL of M9 supplemented with Vitamin B_6_ at **(B)** 1 μM, **(C)** 2.5 μM and **(D)** 5 μM. The starting OD_600_ was 0.08 and OD_600_ reading was measured every 15 minutes during a time course of 40 hours. The hourly OD_600_ data points are reported in the graph. The error bars represent standard deviations from n=3 transformants.

### P5PA are conserved in pathogenic strains of *Pasteurellaceae* family

In order to determine whether this vitamin B_6_ transporter is conserved in microorganisms other than *A. pleuropneumoniae*, we searched for homologues using the P5PA amino acid sequence as a template. A sequence was denoted homologous when it was found to share a minimum of 65% amino acid identity with P5PA. We demonstrated that P5PA homologues are mostly allocated in the family *Pasteurellaceae*. The *Pasteurellaceae* family is predominantly composed of commensal Gram-negative bacteria capable of colonizing the mucosal surfaces of mammals and birds(20, 21). P5PA homologues are exclusively present in the pathogenic strains of the *Pasteurellaceae* family (Fig. 7). The pathogens containing P5PA are closely associated with respiratory diseases and systemic infections in swine (*Actinobacillus*(22)), rabbits, hares, ruminants *(Actinobacillus(23), Mannheimia(24)* and *Pasteurella*(25, 26)), mice (*Muribacter*(27)) and birds (*Pasteurella*(25, 26) and *Gallibacterium*(28)). Moreover, P5PA-encoding bacteria in the genus *Aggregatibacter* are human pathogens and are implicated as the dominant cause of infective endocarditis, brain abscesses (*A. aphrophilus*) and periodontal diseases (*A. actinomycetemcomitans*)(29, 30). Taken together, most of the pathogens in the bacterial family *Pasteurellaceae* possess the P5PA vitamin B_6_ uptake system.

**Figure 7.**
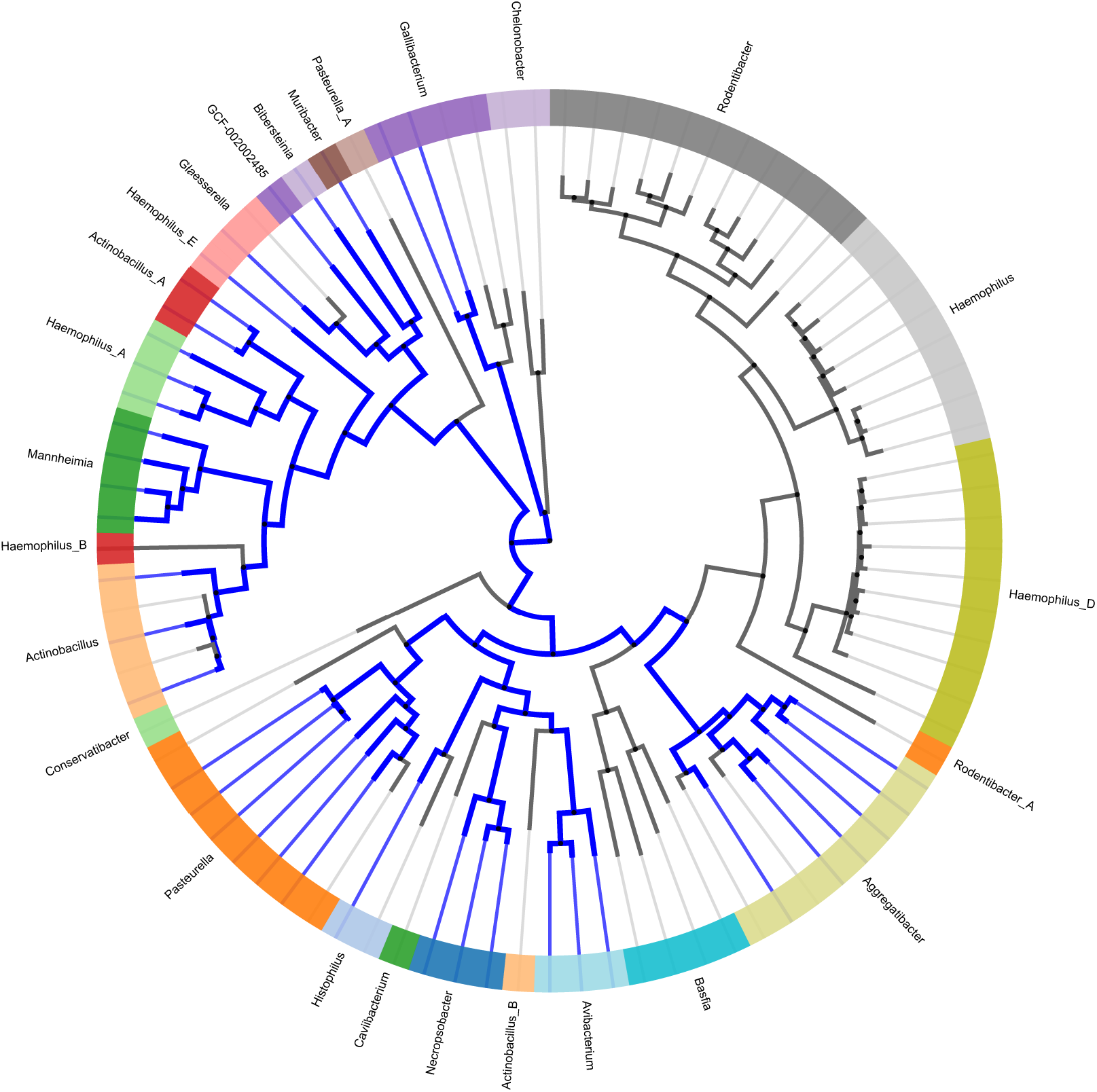
P5PA prevalence in *the Pasteurellaceae*. This tree shows all of the species in the family *Pasteurellaceae*, with the genera labeled and indicated by the colours of the ring. The blue sections of the tree indicate those 38 species containing a P5PA protein as identified by our reciprocal BLAST approach. The tree was produced using AnnoTree (v1.2)(42).

## Discussion

Identified as a homologue of the sugar-phosphate transporter AfuA in *A. pleuropneumoniae*, the functional role of P5PA until now remained controversial. It was hypothesized that P5PA could bind to sugar phosphates just as AfuA, due to the highly similar sequences of these proteins. Another hypothesis was that P5PA could bind iron, which is a limited and essential nutrient required by bacteria for host colonization. In this study, we structurally characterized P5PA and found it to be a type II periplasmic binding protein capable of binding Vitamin B_6_. We confirmed by ITC and LC-MS/MS that the interaction between P5PA and Vitamin B_6_ is highly specific. Moreover, by expressing P5PAB together with the ATPase AfuC from the AfuABC transporter, we were able to rescue the growth of *E. coli* K-12 Δ*pdxB* mutants.

The vitamin B_6_ transport systems in prokaryotes have been poorly characterized. Prior to this study, PdxU1 and PdxU2 were the only proteins predicted to bind a vitamer of vitamin B_6_. This vitamin is called pyridoxine and this prediction was not supported by any experimental data(40). In this study, we illustrate that P5PA transports vitamin B_6_, making it the first reported transporter for the active form of vitamin B_6_ in prokaryotes. P5PA and AfuA share 54% sequence identity as well as structural similarity. However, they are involved in transporting different small nutrient molecules. AfuA plays an important role in transporting G6P, which has a sugar ring and a phosphate moiety, while P5PA is responsible for taking up Vitamin B_6_, which has a pyrimidine ring as well as a phosphate moiety. Comparing the two binding clefts, it becomes clear that the phosphate moieties of G6P and Vitamin B_6_ are both coordinated via hydrogen bonds formed with serine and threonine side chains (Fig.8A, 8B). In the binding pocket of AfuA, Ser9, Arg34, Ser37, Thr60, Ser148, Thr150 and Ser184 surround the phosphate group of G6P. Similarly, the phosphate moiety of P5P is enclosed by Thr9, Thr37, Thr60, Ser146, Ser147 and Thr149 in the binding site of P5PA. However, the ring moieties of G6P and Vitamin B_6_ are stabilized by distinct types of residues (Fig. 8C, 8D). In the binding site of AfuA, the sugar ring is coordinated by three key charged residues, His205, Asp206 and Glu229. It was previously reported that H205A, D206A or E229A mutants of AfuA have undetectable binding affinity for G6P, implying the loss of capacity to sequester G6P(6). However, in the binding cleft of P5PA, the pyrimidine ring of Vitamin B_6_ only interacts with Tyr103 and His204. Most of the residues surrounding His204 are hydrophobic amino acids, including Val203 and Ala205 (Fig. 9), which cannot form electrostatic interactions with the sugar ring of G6P. Furthermore, P5PA does not have highly hydrophilic and charged residues at positions 226-229, which suggests that it cannot coordinate G6P (Fig. 9). It is possible that the substrate specificity of P5PA could be modified to accommodate G6P by making A205D and A228E mutations. In conclusion, P5PA is a highly specific transporter for Vitamin B_6_ and doesn’t possess the capacity to transport G6P due to a lack of hydrophilic and charged residues required to mediate binding of the sugar ring.

**Figure 8.**
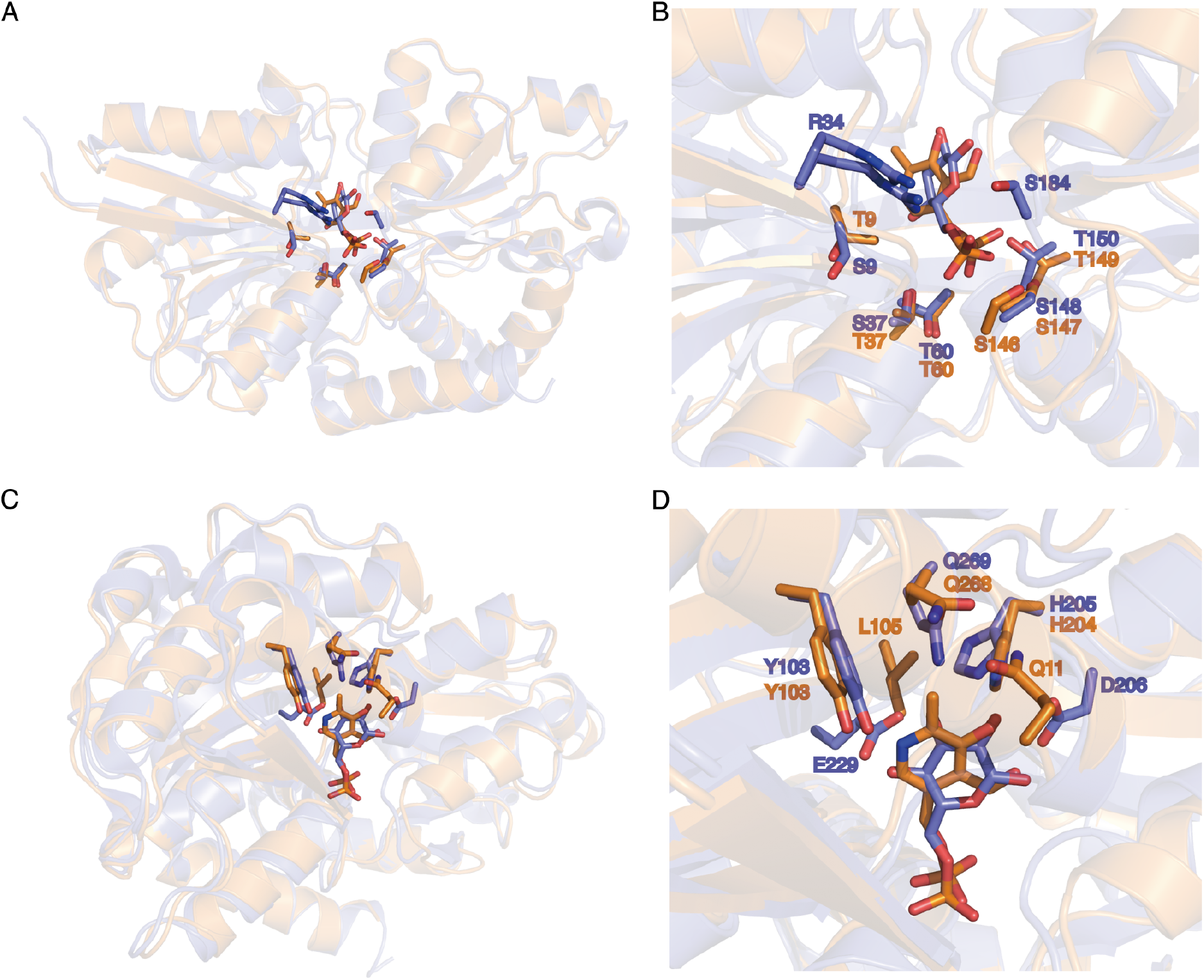
Overlaid structures of P5PA (orange) and AfuA (blue). **(A) (B)**The key residues coordinating the phosphate moieties of G6P and Vitamin B_6_ in AfuA and P5PA respectively. **(C)(D)** The residues stabilizing the ring moieties of G6P and Vitamin B_6_ in AfuA and P5PA respectively

**Figure 9.**
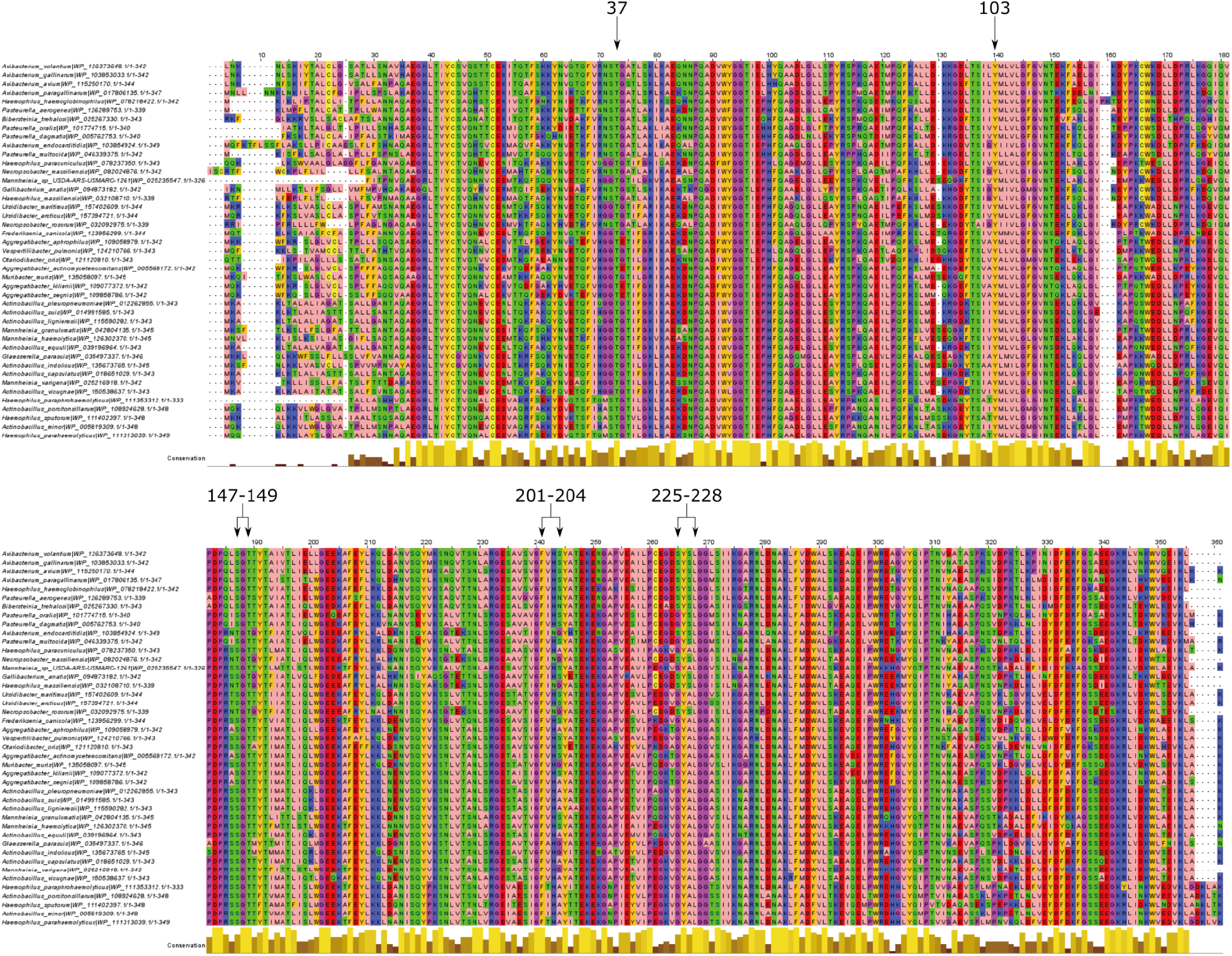
Multiple sequence alignment of putative P5PA sequences. Residues are coloured according to their chemical properties, and the black arrows indicate regions from the A. pleuropneumoniae protein P5PA sequence discussed in the text. The alignment was generated using the G-INS-i algorithm from MAFFT (v7.450)(43), and has been visualized using JalView (v2.11.0)(44).

Vitamin B_6_ functions as an essential cofactor for a variety of cellular catalytic reactions and nutrient metabolisms including amino acids, fatty acids, and carbohydrates(31). It is also implicated in regulating cell stress and oxidants(32, 33). In most microorganisms, approximately 1.5% of genes encode Vitamin B_6_-dependent enzymes which are responsible for catalyzing more than 140 biochemical reactions(31, 34). Thus, acquiring Vitamin B_6_ is critical for prokaryote survival. Bacteria have two independent *de novo* pathways to synthesize Vitamin B_6_. Most γ-proteobacteria, including *E. coli*, exploit the deoxyxylulose 5-phosphate (DXP)-dependent pathway where D-erythrose 4-phosphate undergoes a series of biochemical reactions, the product of which then interacts with DXP to form the P5P precursor, pyridoxine-5’-phosphate (PNP). This reaction is catalyzed by two key enzymes, PdxA and PdxJ (32, 33). PNP is then converted to pyridoxial-5-phosphate (P5P, active form of Vitamin B_6_) via a salvage pathway. On the other hand, the DXP-independent pathway is found in the majority of bacteria, and also in archaea, fungi, and plants. In this pathway, Vitamin B_6_ is derived from glutamine, ribose 5-phosphate and GAP with the help of two critical enzymes, PdxS (also referred to as Pdx1) and PdxT (also referred to as Pdx2)(32, 35). The DXP-independent pathway was recently characterized in *A. pleuropneumoniae* by Xie et al(36). When the authors disrupted the *pdx*S and *pdx*T genes in *A. pleuropneumoniae*, the resulting mutants showed a significant decrease in growth in the absence of Vitamin B_6_. However, supplementation with 10 μM Vitamin B_6_ rescued growth(36). These data agree with our observation that there is no significant difference in growth between wild-type *E. coli* K-12 and the Δ*pdxB* mutant at 5 μM and 10 μM Vitamin B_6_. This may indicate that, at a high concentration (≥5 μM), exogenous Vitamin B_6_ can diffuse into bacterial cells via passive transport. In fact, previous studies have illustrated that other dephosphorylated vitamers of vitamin B_6_ enter *E.coli* K-12 and lactic acid bacteria by facilitated diffusion(37, 38).

The Vitamin B_6_ concentration in swine plasma is maintained at a low level of 36-45 nM (39), which suggests that passive movement of Vitamin B_6_ across the bacterial membrane is inefficient in the swine host. Hence, possessing a nutrient transporter specific for Vitamin B_6_ increases the chance of successful colonization by *A. pleuropneumoniae* when its vitamin B_6_ biosynthesis pathway is dysfunctional due to deficiencies in starting materials for Vitamin B_6_ synthesis. This hypothesis was supported in the present study by the observed effect of expressing the P5PAB together with AfuC in *ΔpdxB* mutants. Expression of the *p5pA*B+*afu*C operon enabled Δ*pdxB* mutants to overcome the growth deficiency observed at low concentrations of Vitamin B_6_. Another possible reason for expressing a Vitamin B_6_-specific transporter is that nutrient transport is favored over nutrient biosynthesis by microorganisms, since the former consumes less energy (40). In addition to vitamin B_6_, other B-type vitamins are also subject to extracellular uptake by transporters in the ATP binding cassette (ABC) transporter family. For example, the important cofactor vitamin B12 can be taken up from the extracellular environment in *E. coli* by the ABC importer BtuCDF and synthesized by the enzymes encoded in the *cod* operon(40, 41). Overcoming nutrient scarcity using these specific transport systems permits bacteria to strive in a variety of inhospitable environments but also to outcompete neighboring bacteria for key nutrients including vitamins.

## Acknowledgements

We thank members of APS at the NE-CAT beamlines 24-ID-E and 24-ID-C and CLS beamline staff at CMCF-08ID-1 for assistance with data collection. We also thank members of the Moraes laboratory for valuable discussion. Dr. Karen Maxwell provided the Keio Collection *E.coli* strains. We would like to acknowledge the Analytical Facility for Bioactive Molecules (AFBM) of the Centre for the Study of Complex Childhood Diseases (CSCCD) at The Hospital for Sick Children, Toronto, Ontario, especially Ashley St-Pierre for her assistance with LC/MS/MS.

## Supplementary Information

There are 2 supplementary figures and 1 supplementary table associated with this manuscript.

**Supplementary Table1.**
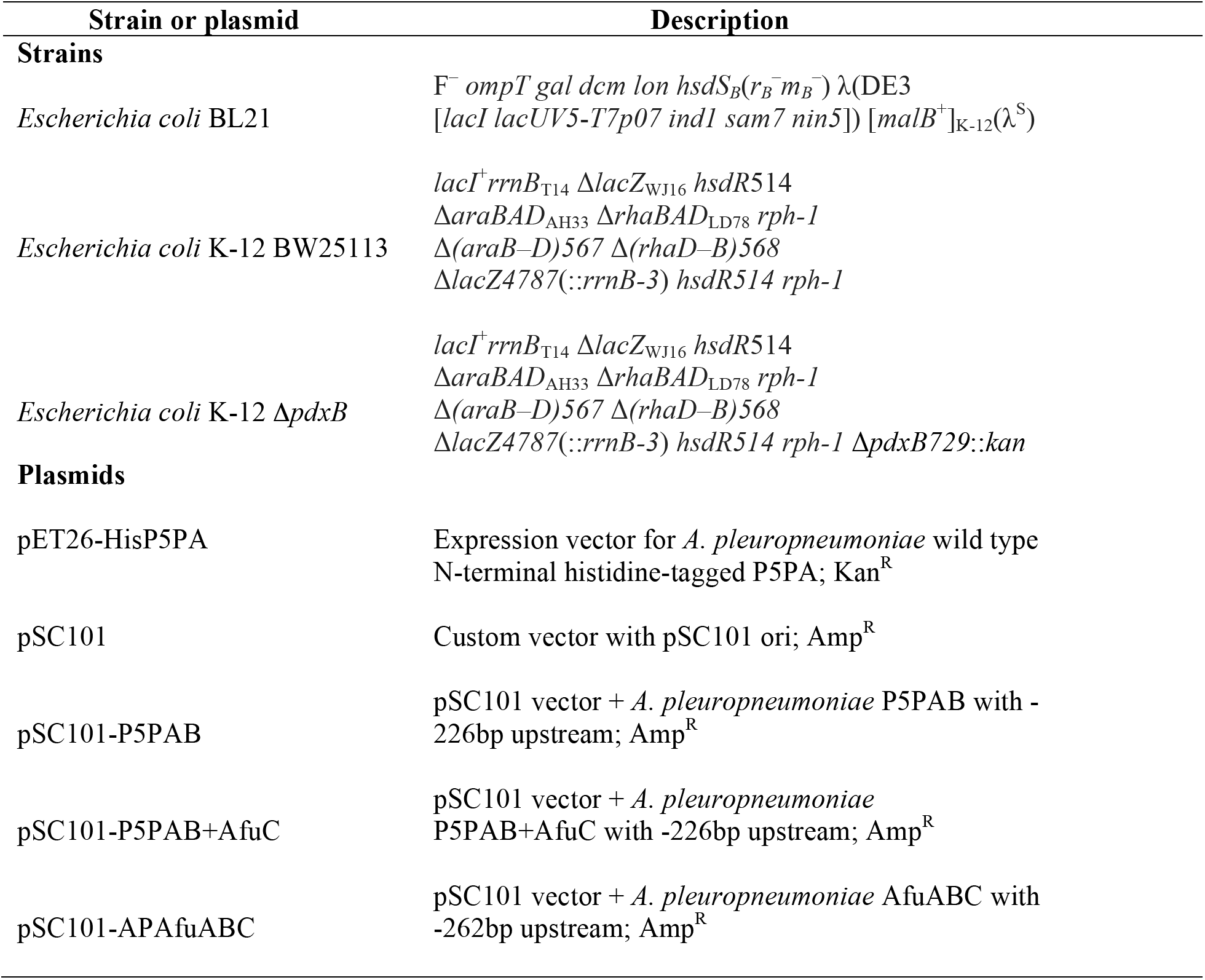
Strains and plasmids used in this study

**Supplementary Figure 1.**
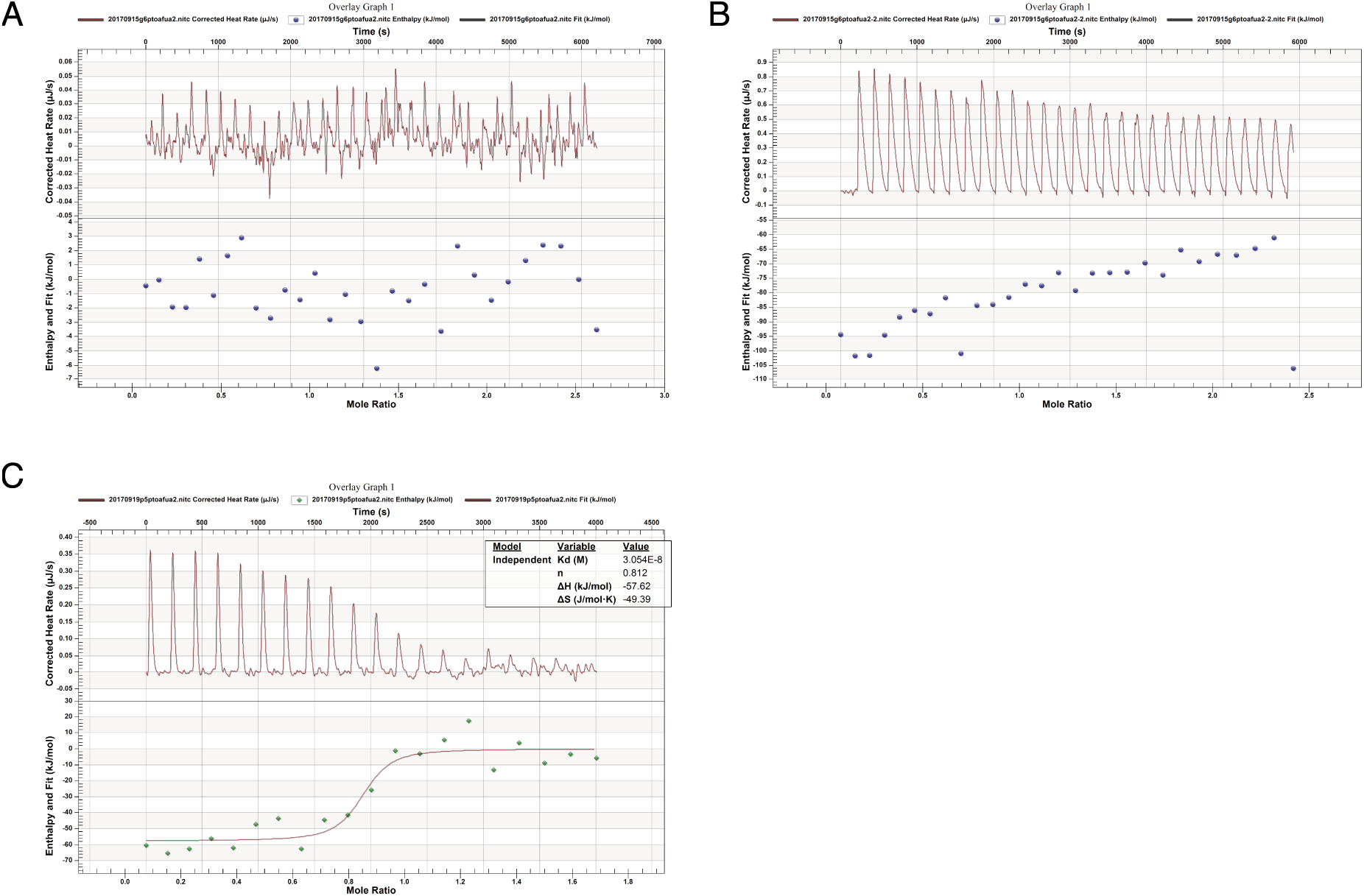
G6P does not bind to P5PA. **(A)** Biological replicate n=1 **(B)** Biological replicate n=2 ITC diagram of the interaction between AfuA2 and G6P. 45 μM of P5PA was titrated with 300 μM of G6P which is measured by ITC-TA. **(C)** G6P-titrated P5PA was collected from the sample cell in ITC instrument and separated from G6P by gel filtration in S75 column. It was titrated against 300 μM Vitamin B_6_ to obtain Kd=35 nM implying the used AfuA2 was active and functional.

**Supplementary Figure 2.**
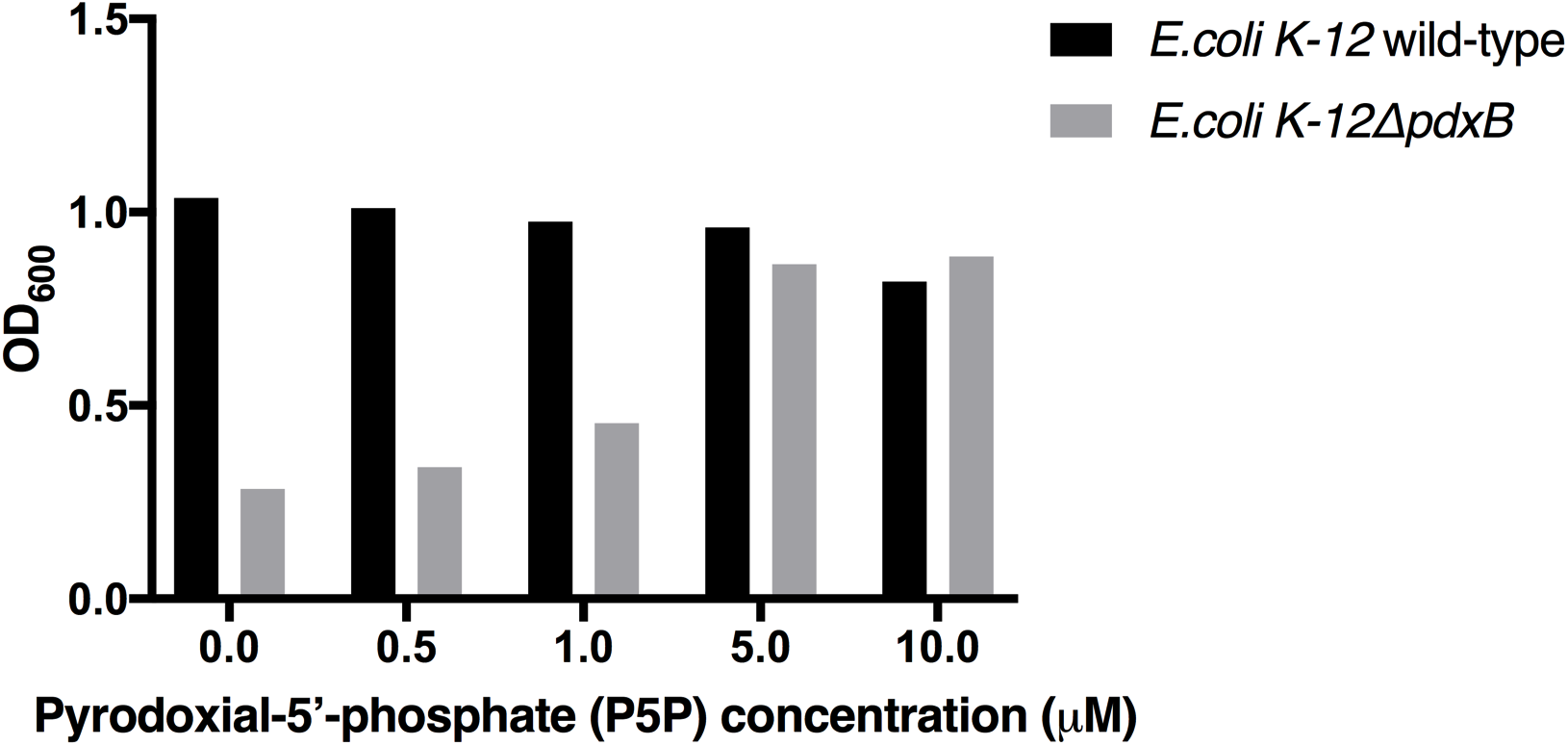
*E. coli* K-12 Δ*pdxB* mutant has a growth deficiency in M9 supplemented with Vitamin B_6_ (Pyrodoxial-5’-phosphate-P5P) at 0.5 μM and 1 μM. With starting OD_600_=0.05, *E. coli* K-12 wild-type and Δ*pdxB* mutant were incubated in 5 mL of M9 supplemented with Vitamin B_6_ at a series concentration of 0.5 μM, 1 μM, 5 μM and 10 μM. OD_600_ was measured after a 23 hour incubation at 37°C.

